# *Plasmodium berghei* kinesin-5 associates with the spindle apparatus during cell division and is important for efficient production of infectious sporozoites

**DOI:** 10.1101/2020.07.03.186031

**Authors:** Mohammad Zeeshan, Declan Brady, Rebecca R. Stanway, Carolyn A. Moores, Anthony A. Holder, Rita Tewari

## Abstract

Kinesin-5 motors play essential roles in spindle apparatus assembly during cell division, by generating forces to establish and maintain the spindle bipolarity essential for proper chromosome segregation. Kinesin-5 is largely conserved structurally and functionally in model eukaryotes, but its role is unknown in the *Plasmodium* parasite, an evolutionarily divergent organism with several atypical features of both mitotic and meiotic cell division. We have investigated the function and subcellular location of kinesin-5 during cell division throughout the *Plasmodium berghei* life cycle. Deletion of *kinesin-5* had little visible effect at any proliferative stage except sporozoite production in oocysts, resulting in a significant decrease in the number of motile sporozoites in mosquito salivary glands, which were able to infect a new vertebrate host. Live-cell imaging showed kinesin-5-GFP located on the spindle and at spindle poles during both atypical mitosis and meiosis. Fixed-cell immunofluorescence assays revealed kinesin-5 co-localized with α-tubulin and centrin-2 and a partial overlap with kinetochore marker NDC80 during early blood stage schizogony. Dual-colour live-cell imaging showed that kinesin-5 is closely associated with NDC80 during male gametogony, but not with kinesin-8B, a marker of the basal body and axonemes of the forming flagella. Treatment of gametocytes with microtubule-specific inhibitors confirmed kinesin-5 association with nuclear spindles and not cytoplasmic axonemal microtubules. Altogether, our results demonstrate that kinesin-5 is associated with the spindle apparatus, expressed in proliferating parasite stages, and important for efficient production of infectious sporozoites.

## Introduction

Kinesin-5 proteins are a family of molecular motors that is structurally and functionally conserved throughout eukaryotes (Mann and Wadsworth, 2019; Waitzman and Rice, 2014; Wojcik et al., 2013). They are involved in spindle pole separation and are considered essential for mitosis in the vast majority of eukaryotes (Bannigan et al., 2007; Ferenz et al., 2010), except *Caenorhabditis elegans* (Bishop et al., 2005), *Dictyostelium discoideum* (Tikhonenko et al., 2008) and *Candida albicans* (Shoukat et al., 2019). Kinesin-5 contains an N-terminal kinesin motor domain, a central stalk and a C-terminal tail domain (Wojcik et al., 2013), and forms a bipolar homotetramer that cross-bridges and slides on parallel and anti-parallel microtubules (MTs) (Kapitein et al., 2008). The motor domain binds microtubules (MTs) and hydrolyses ATP, which are conserved functions, while the tail region can also bind MTs, regulate motor activity (Bodrug et al., 2020) and help localization during mitosis (Bodrug et al., 2020; Weinger et al., 2011). The stalk contains a coiled-coil region and neck linker that promote oligomerization and direction of movement, respectively (Hesse et al., 2013). The kinesin-5 motor domain is conserved across eukaryotes including *Plasmodium* **(Fig. S1A),** but the tail region is highly variable except for a short, conserved region called the BimC box, which contains consensus sites for phosphorylation by cyclin dependent kinase 1 (Cdk1) in most eukaryotes (Bishop et al., 2005; Chee and Haase, 2010; Sharp et al., 1999).

Kinesin-5 is located at spindle MTs and spindle poles during cell division and is distributed diffusely in the cytoplasm during interphase in most eukaryotic cells (Ferenz et al., 2010; Waitzman and Rice, 2014). The essential roles of kinesin-5 in spindle assembly and spindle pole separation can be blocked by specific inhibitors or antibodies, resulting in collapse of bipolar spindles into monopoles (Ferenz et al., 2010; Kapoor et al., 2000; Mann and Wadsworth, 2019; Sharp et al., 1999). Kinesin-5 is required for maintenance of a bipolar spindle in fungi, Xenopus and Drosophila (Hoyt et al., 1992; Kapoor et al., 2000; Sharp et al., 1999). In budding yeast the kinesin-5 proteins, Cin8 and Kip1, are also present at the kinetochores and help in chromosome alignment during metaphase (Tytell and Sorger, 2006).

Malaria is a deadly vector-borne infectious disease, caused by a unicellular protozoan parasite of the genus *Plasmodium*, which infects many vertebrate hosts including humans, and is transmitted by female Anopheles mosquitoes (WHO, 2019). During the complex life cycle two unique phases of atypical closed mitotic division occur. The first type of mitosis occurs during asexual proliferation, with multiple asynchronous nuclear divisions producing up to 32 nuclei (during schizogony in the blood of the vertebrate host) or more than 1000 nuclei (during schizogony in the liver of the vertebrate host and sporogony in the mosquito gut), with cytokinesis occurring only after nuclear division is complete, to produce haploid progeny cells (Francia and Striepen, 2014; Gerald et al., 2011; Sinden, 1991; Sinden et al., 1976). It is important to note that the final round of nuclear division is asynchronous, in contrast to what was thought previously (Rudlaff et al., 2020). The second type of mitosis is during male gametogony (part of the sexual stage in the mosquito gut) where there are three rapid rounds of DNA replication from 1N to 8N within 10 to 15 minutes, followed by karyokinesis and cytokinesis to produce 8 flagellate haploid male gametes (Sinden et al., 1976; Zeeshan et al., 2019a). The first phase of meiotic division occurs following fertilization as the zygote differentiates into a motile ookinete in the mosquito gut. The 2N genome is duplicated and recombination occurs (Sinden, 1991); then the final reductive division likely occurs in the oocyst that is formed following ookinete penetration of the mosquito gut wall, leading to the formation of haploid sporozoites. Thus, *Plasmodium* has unusual ways to divide and survive under different physiological conditions in different hosts. As cell division in *Plasmodium* is atypical, so the molecules regulating cell division are also divergent from those of higher eukaryotes (Guttery et al., 2014; Roques et al., 2015; Tewari et al., 2010; Wall et al., 2018). The large family of kinesins includes molecular motors that are essential for several processes during cell division (Wordeman, 2010; Yount et al., 2015), and in *Plasmodium berghei* there are nine kinesin genes, including two kinesin-8 genes that are important in cell division and male gamete formation (Zeeshan et al., 2019a; Zeeshan et al., 2019b). There is a single *Plasmodium* kinesin-5, and since this protein plays an important role during cell proliferation in many eukaryotes, and is considered as a strong target for the development of therapeutics against many diseases including malaria (Liu et al., 2014), we decided to study its role during cell division in the malaria parasite.

In the present study we examined the spatiotemporal dynamics and functional role of kinesin-5 during the proliferative stages of *P. berghei*. Unlike kinesin-5 in many other eukaryotes, *P. berghei* (Pb) kinesin-5 is dispensable for parasite growth and proliferation, although deletion of the gene results in a remarkable decrease in the number of mosquito salivary gland sporozoites. Live cell imaging shows that kinesin-5 is located both on the spindle and at spindle poles during mitotic and meiotic divisions in the parasite life cycle, co-localizing with centrosome marker centrin and kinetochore marker NDC80. Treatment of gametocytes with microtubule inhibitors confirmed the nuclear localization and association with spindle microtubules of kinesin-5 during male gametogony. Altogether, our results show that kinesin-5 is associated with the spindle apparatus, which can be inhibited by microtubule inhibitors, and that the protein is important for efficient production of infectious sporozoites.

## Results

### Pbkinesin-5 is expressed at multiple proliferative stages in the parasite life cycle

To quantify the expression of kinesin-5 at different stages of the parasite life cycle, we isolated RNA and performed qRT-PCR. *Kinesin-5* is expressed constitutively throughout the blood and mosquito stages of parasite development, with the highest level in gametocytes, followed by schizonts and ookinetes (**Fig. 1A**).

**Fig. 1.**
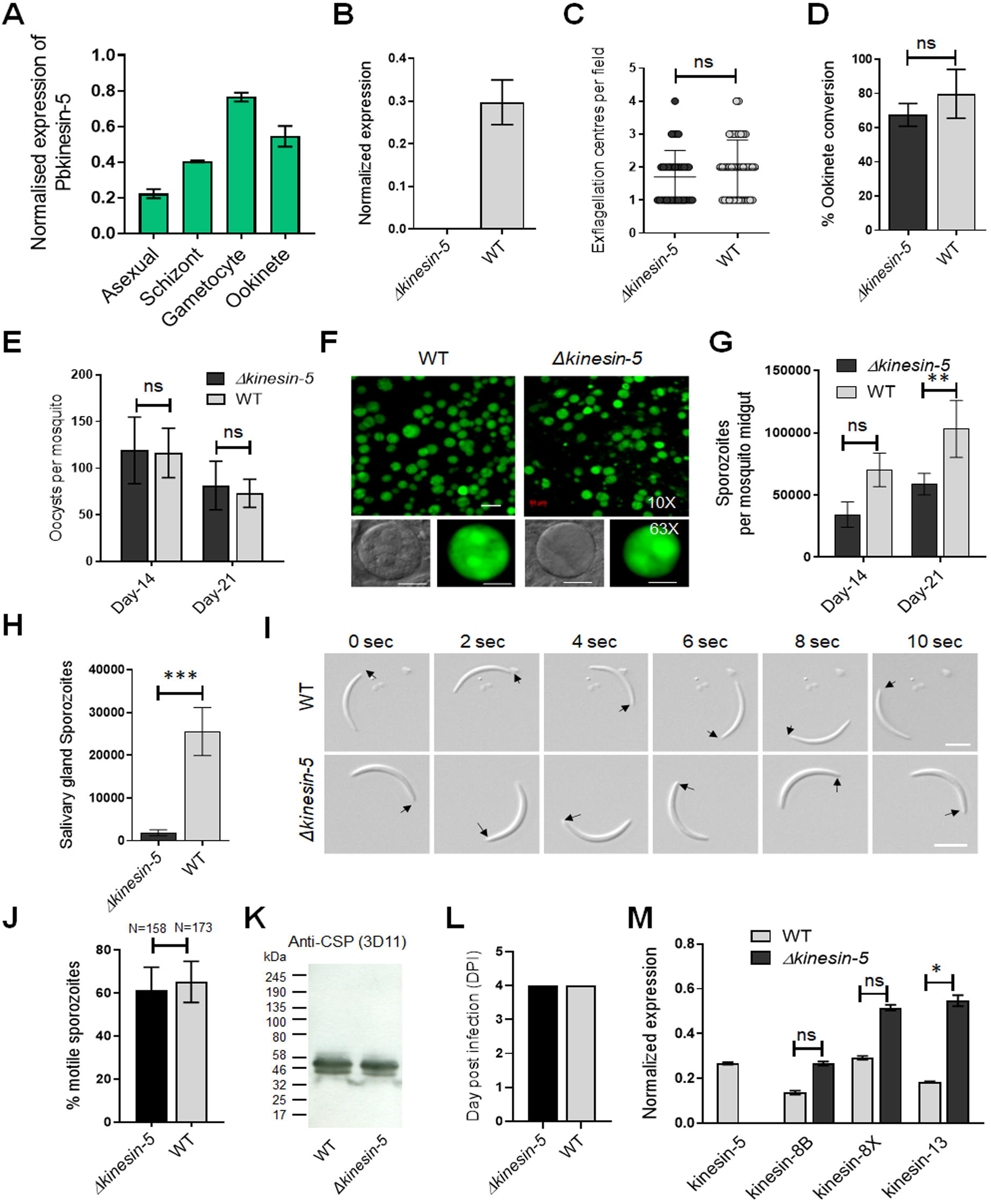
The *Pb*kinesin-5 gene is dispensable for parasite transmission but has a role in sporozoite development. **(A)** Transcript levels of kinesin-5 revealed by qRT-PCR, normalised against two endogenous control genes, arginine-tRNA synthetase and hsp70. Each bar is the mean of three biological replicates ± SD. **(B)** qRT-PCR analysis of *kinesin-5* transcription in *Δkinesin-5* and WT-GFP parasites, showing the complete depletion of *kinesin-5*. Each bar is the mean of three biological replicates ± SD. **(C)** Male gametogony (exflagellation) of *Δkinesin-5* line (black bar) and WT-GFP line (grey bar) measured as the number of exflagellation centres per field. Mean ± SD; n=6 independent experiments. **(D)** Ookinete conversion as a percentage for *Δkinesin-5* (black bar) and WT-GFP (grey bar) parasites. Ookinetes were identified using 13.1 antibody as a surface marker and defined as those cells that differentiated successfully into elongated ‘banana shaped’ ookinetes. Mean ± SD; n=6 independent experiments. **(E)** Total number of GFP-positive oocysts per infected mosquito in *Δkinesin-5* (black bar) and WT-GFP (grey bar) parasites at 14- and 21-days post infection (dpi). Mean ± SD; n=5 independent experiments. **(F)** Mosquito mid guts at 10x and 63x magnification showing oocysts of *Δkinesin-5* and WT-GFP lines at 14 dpi. Scale bar = 50 μm in 10x and 20 μm in 63x. **(G)** Total number of sporozoites in oocysts of *Δkinesin-5* (black bar) and WT-GFP (grey bar) parasites at 14 and 21 dpi. Mean ± SD; n=4 independent experiments. **(H)** Total number of sporozoites in salivary glands of *Δkinesin-5* (black bar) and WT-GFP (grey bar) parasites. Bar diagram shows mean ± SD; n=4 independent experiments. **(I)** Differential interference contrast (DIC) time-lapse image sequences showing motile *Δkinesin-5* and WT-GFP sporozoites isolated from salivary glands. Arrow indicates apical end of sporozoites. Scale bar = 5 μm. **(J)** Quantitative data for motile sporozoites from salivary glands for *Δkinesin-5* and WT-GFP based on two independent experiments. **(K)** Western blot analysis of WT-GFP and *Δkinesin-5* parasites. Lysates from midgut sporozoites were probed using a monoclonal antibody specific for circumsporozoite protein (CSP)-repeat region (mAb 3D11). Molecular mass markers (kDa) are shown on the left of the gel. **(L)** Bite back experiments show successful transmission of *Δkinesin-5* parasites (black bar) from mosquito to mice, similar to WT-GFP parasites (grey bar). Mean ± SD; n= 3 independent experiments. **(M)** qRT-PCR analysis of other Pbkinesin genes comparing transcript levels in WT-GFP and *Δkinesin-5* parasites. Error bar = ±SD; n=3. Unpaired t-test was performed for statistical analysis. *p<0.05, **p<0.01, ***p<0.001, ns=non-significant.

### Pbkinesin-5 is important for efficient production of infectious sporozoites

The *Plasmodium* life cycle has two unusual mitotic processes that occur during schizogony/sporogony and male gametogony, and a single meiotic stage during zygote to ookinete transformation (Zeeshan et al., 2020b). To examine any functional role of kinesin-5 during these processes, we deleted the gene from the *P. berghei* genome using a double crossover homologous recombination strategy in a parasite line constitutively expressing green fluorescent protein (GFP) at all stages of the parasite life cycle (**Fig. S1B**) (Janse et al., 2006). Diagnostic PCR, to show successful integration of the targeting construct at the *kinesin-5* locus (**Fig. S1C**), and quantitative real time PCR (qRT-PCR), to show lack of *kinesin-5* expression in gametocytes, confirmed the complete deletion of the *kinesin-5* gene (**Fig. 1B**). Successful creation of this transgenic parasite indicated that the gene is not essential for mitosis during asexual blood stage schizogony. Further phenotypic analysis of the *Δkinesin-5* parasite was carried out at other stages of the life cycle, comparing the parental parasite (WT-GFP) with two independent gene-knockout parasite clones (clones 3 and 5) generated by two independent transfections. Both gene-knockout clones had the same phenotype and data presented here are the combined results from both clones. Since *Δkinesin-5* parasites underwent asexual blood stage development, exhibiting no change in morphology, number of progeny merozoites or parasitemia, an essential role for kinesin-5 is unlikely during these stages that cause the disease in the mammalian host.

The transgenic parasites also produced gametocytes in mice, and therefore next we analysed male and female gametocyte differentiation following activation in the exflagellation/ookinete medium that mimics the mosquito gut environment. Male and female gametes emerge from the infected erythrocyte, and in the case of male gamete development this is preceded by three rapid rounds of genome duplication, resulting in eight flagellate male gametes (Sinden et al., 1976; Zeeshan et al., 2019a). There was no defect in male gamete exflagellation for either of the *Δkinesin-5* parasite clones, with the same exflagellation frequency as WT-GFP parasites (**Fig. 1C**). Fertilization, zygote formation and ookinete differentiation, when meiosis occurs (Sinden, 1991), were not significantly different in *Δkinesin-5* parasites from these processes in parental parasites (**Fig. 1D**).

To investigate the role of kinesin-5 in oocyst development and sporogony, *Anopheles stephensi* mosquitoes were fed on mice infected with *Δkinesin-5* parasites and WT-GFP parasites as a control. The number of GFP-positive oocysts on the mosquito gut wall was counted on days 14 and 21 post-infection. There was no significant difference in the number of *Δkinesin-5* and WT-GFP oocysts (**Fig. 1E**), and the size of the oocysts was similar for both parasites (**Fig. 1F**). However, we observed a 40 to 50% decrease in the number of sporozoites in each oocyst at days 14 and 21 post-infection in *Δkinesin-5* parasites compared to WT-GFP parasites (**Fig. 1G**) indicating the important role of kinesin-5 during sporogony. In comparison to the WT-GFP parasite, a significant decrease in the number of *Δkinesin-5* sporozoites in salivary glands was also observed (**Fig. 1H**), but the shape, size, and motility of these sporozoites were indistinguishable from WT-GFP parasites (**Fig. 1I, J**). Furthermore, western blot analysis of circumsporozoite protein (CSP) showed that proteolytic processing of CSP in *Δkinesin-5* sporozoites, an indicator of normal sporozoite maturation, was also not affected (**Fig. 1K**). The infected mosquitoes were used for bite back experiments to ascertain the infectivity of *Δkinesin-5* sporozoites in mice; a blood stage infection was observed after 4 days with both *Δkinesin-5* and WT-GFP sporozoites (**Fig. 1L**).

The loss of kinesin-5 may have been compensated for by the over-expression of another member of the kinesin family, therefore we analysed the transcript level of kinesin-8B, kinesin-8X and kinesin-13 in gametocytes of *Δkinesin-5* parasites. These kinesins are highly expressed in gametocytes and have important role during male gametogony (Zeeshan et al., 2019a; Zeeshan et al., 2019b). We found that the kinesin-13 transcript level was significantly upregulated (**Fig. 1M**), suggesting that higher levels of kinesin-13 may - in some way - compensate for the kinesin-5 deletion.

### Pbkinesin-5 is located at the spindle apparatus during mitotic stages of asexual blood stage schizogony

To examine expression at the protein level and study the real-time dynamic location of kinesin-5 during cell division, we generated a kinesin-5-GFP transgenic *P. berghei* line expressing kinesin-5 with a C-terminal GFP tag, by inserting an in-frame *gfp* coding sequence at the 3’ end of the endogenous *kinesin-5* locus using single homologous recombination (**Fig. S2A**). Successful insertion was confirmed by diagnostic PCR (**Fig. S2B**). Western blot analysis of a schizont protein extract using an anti-GFP antibody revealed kinesin-5-GFP protein at the expected size of 198 kDa compared to the 29 kDa GFP (**Fig. S2C**). This kinesin-5-GFP transgenic line was used to examine the spatiotemporal profile of kinesin-5-GFP protein expression and location by live cell imaging during the whole parasite life cycle, initially during asexual blood stage development in erythrocytes.

After haploid merozoite invasion of an erythrocyte, the initial ring and trophozoite stages are followed by schizogony, which results in the formation of further merozoites that invade fresh erythrocytes. During schizogony there are several independent asynchronous rounds of closed mitosis to produce a multi-nucleate coenocyte, followed by cytokinesis and egress of mature merozoites from the infected erythrocyte. Kinesin-5 expression was not detected in the ring and early trophozoite stages that are considered as interphase or G_0_ in the cell cycle (Arnot and Gull, 1998). A very low and diffuse expression of kinesin-5 throughout the cytoplasm was observed in older trophozoites, a stage similar to G1 phase in higher eukaryotes (Arnot and Gull, 1998; Arnot et al., 2011) when preparation for DNA replication begins. Late trophozoites mark the transition into early S phase when DNA synthesis starts (Arnot et al., 2011) and schizogony is marked by the presence of three or more nuclei (Rudlaff et al., 2020). Kinesin-5 was observed as strong foci adjacent to the Hoechst-stained DNA in early schizonts when nuclear division had commenced, and representing the first M phase of the cell cycle (Arnot and Gull, 1998; Arnot et al., 2011) (**Fig. 2A**). Each kinesin-5-GFP focus elongated and split into two foci that migrated away from each other, remaining adjacent to the nuclear DNA that then separated into two nuclear masses (**Fig. 2A**). Alternating repeated S/M phases followed the division of individual nuclei, accompanied by repeated elongation and duplication into multiple points of these kinesin-5-GFP foci, showing the asynchronous pattern of nuclear division. Following completion of schizogony, kinesin-5 expression was almost undetectable in merozoites **(Fig. 2A).**

**Fig. 2.**
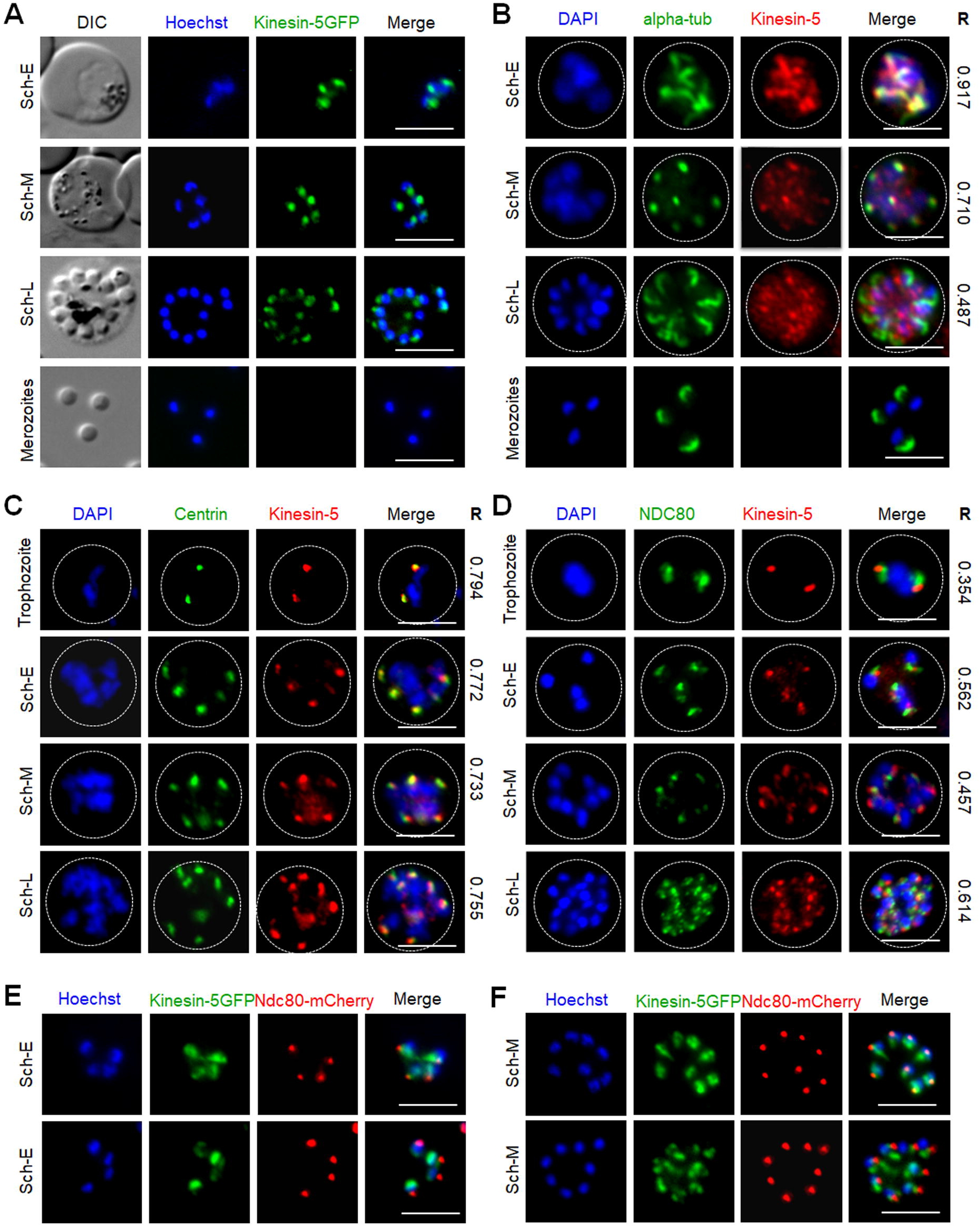
Localization of Pbkinesin-5 on the spindle and spindle pole during asexual blood stage schizogony and its association with other cell division markers. **(A)** Live imaging of *Pb*kinesin-5-GFP (Green) during schizogony within a host erythrocyte, showing its location on a putative MT organizing centre (MTOC) and mitotic spindle during early dividing stages. The protein is diffuse or absent in mature merozoites. **(B-D)** Indirect immunofluorescence assays (IFA) showing the location of *Pbk*inesin-5 (red) in relation to α-tubulin (green, B), centrin (green, C) and NDC80 (green, D). Dotted lines represent the red blood cell membrane. **(E, F)** Live imaging showing the location of kinesin-5–GFP (green) in relation to NDC80-mCherry (red). DIC: Differential interference contrast. Sch-E (Early schizont), Sch-M (Middle schizont), Sch-L (Late schizont), Scale bar = 5 μm, R (Pearson’s coefficient for co-localization).

To compare the location of kinesin-5 with that of other mitotic protein markers, including α-tubulin (spindle MTs), centrin-2 (putative centrosome/spindle pole body [SPB]/ MT organising centre [MTOC]) and NDC80 (kinetochores), we used indirect immunofluorescence (IFA)-based co-localization assays with anti-GFP antibodies and other antibodies specific for the marker proteins. We observed co-localization of kinesin-5 with α-tubulin at the early stages of schizogony, both at spindle MTs and the putative MTOC with a Pearson’s colocalization coefficient (R) of more than 0.7 (**Fig. 2B**). Similarly, kinesin-5 co-localised with centrin-2 with a Pearson’s colocalization coefficient (R) of more than 0.7, confirming its location close to or at the putative MTOC (**Fig. 2C)**. Using anti-GFP with anti-NDC80 antibodies revealed that kinesin-5-GFP is located in close proximity to, and partially overlapping, the kinetochores with a Pearson’s colocalization coefficient (R) of less than 0.7 (**Fig. 2D)**; this location was confirmed by live cell imaging with a dual colour parasite line expressing kinesin-5-GFP and NDC80-mCherry (**Fig. 2E, F)**.

### Spatiotemporal dynamics of Pbkinesin-5 reveal its location on the spindle apparatus during male gametogony

In order to study the dynamics of kinesin-5 during the rapid genome replication in male gametogony we examined its expression by live-cell imaging during the 15-minute period following activation of male gametocytes. Both male and female gametocytes express kinesin-5, with a diffuse nuclear location. At the start of male gametogony, kinesin-5 accumulated at one end of the nucleus at a single focal point 1-minute post-activation (mpa) (**Fig. 3A**). By 2 mpa, this focal point extended to form a bridge across one side of the nucleus, followed by the separation of the two halves of the bridge to produce shorter linear rods that then contracted to two clear single foci by 3 mpa (**Fig. 3A)**. This process repeated twice, resulting in 8 discrete kinesin-5-GFP foci. These discrete kinesin-5 foci then dispersed just before cytokinesis and exflagellation and the protein remained diffused in the nucleus of the remnant gametocyte (**Fig. 3A)**. A schematic diagram for this process is shown in the upper panel of **Fig. 3A**.

**Fig. 3.**
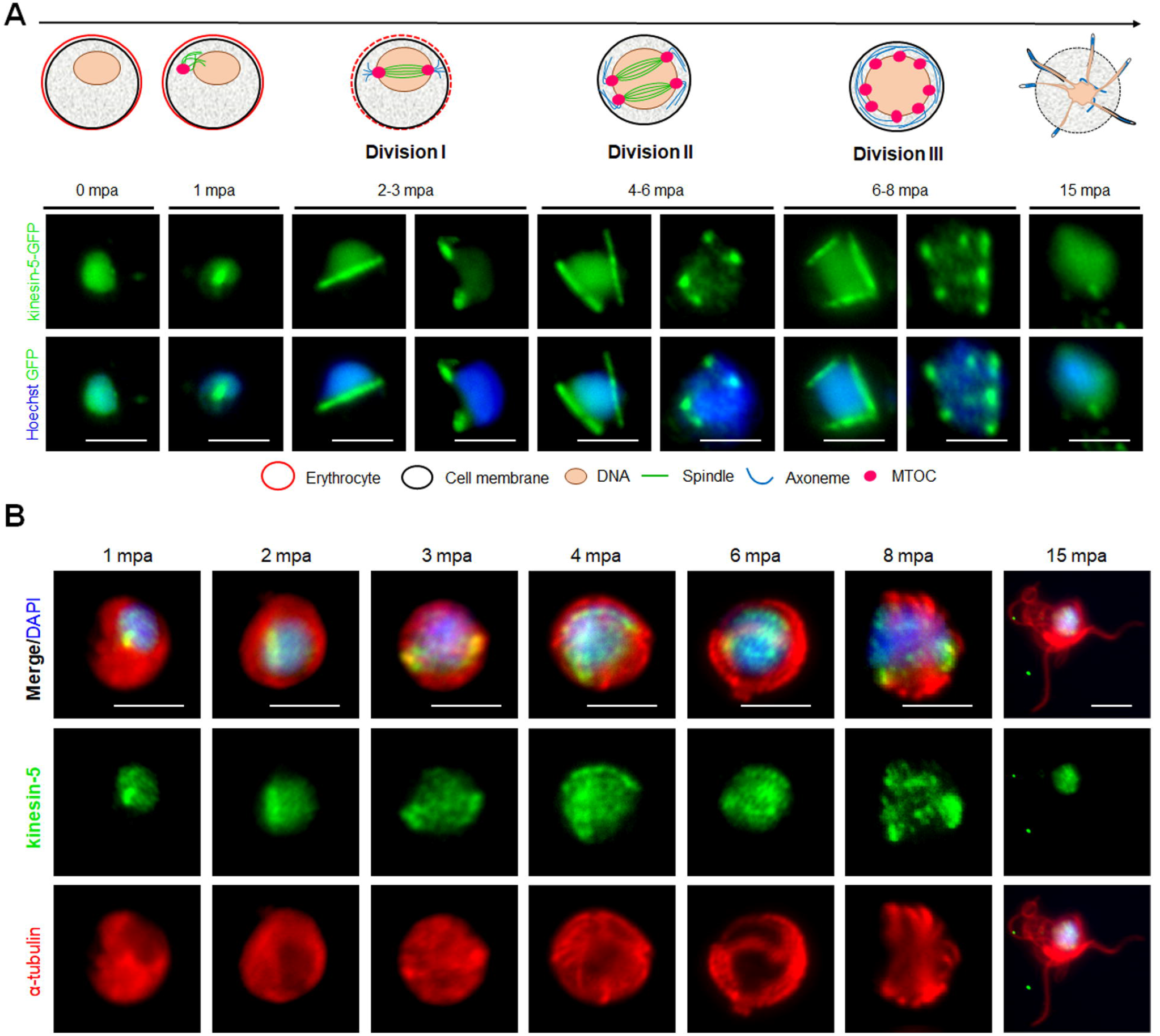
The location of Pbkinesin-5 and its association with MTs during male gametogony. **(A)** Live-cell imaging of Pbkinesin-5-GFP (Green) during male gametogony showing an initial location on a putative MT organizing centre (MTOC) just after activation, and then on spindles and spindle poles in the later three mitotic stages. The schematic shows the principle stages of male gametogony. **(B)** IFA showing location of kinesin-5 (green) and α-tubulin (red) in male gametocytes at 1-, 2-, 3-, 4-, 6-, 8- and 15-min post activation (mpa). Scale bar = 5 μm.

To study the association of kinesin-5 with the mitotic spindle we used immunofluorescence-based co-localization assays with anti-GFP antibodies to stain kinesin-5, and anti-α-tubulin antibodies for MT staining. This analysis showed clear co-localisation of kinesin-5-GFP during early stages (1-3 mpa) of male gametogony with the spindle MT, both on the bridge-like structure and the foci, representing the spindle and spindle pole body, respectively (**Fig. 3B**). Because the anti-α-tubulin antibodies recognise both spindle (nuclear) and axonemal (cytoplasmic) MT, spindle MTs are not distinctly visualised during later stages (6-8 mpa) when axoneme assembly is nearly complete. However, it is very clear that the pattern of kinesin-5 staining does not correspond with axonemal MT. Furthermore, kinesin-5 was not present in mature male gametes following their egress **(Fig. 3A, B).**

To investigate further whether the location of kinesin-5 is cytoplasmic or nuclear, the kinesin-5-GFP parasite line was genetically crossed with the NDC80-mCherry kinetochore marker line (Zeeshan et al., 2020b) and kinesin-8B-mCherry cytoplasmic axoneme marker line (Zeeshan et al., 2019a), and the crosses were used for live cell imaging of both markers to establish their spatiotemporal relationship. We found that both kinesin-5 and NDC80 were located next to the nuclear DNA, with co-localization on both spindle and spindle poles during different stages of male gametogony **(Fig. 4A, B)**. In contrast, kinesin-5 did not co-localise with kinesin-8B which is located on the cytoplasmic basal bodies in early stages of male gametogony and later distributes across the axonemes (**Fig. 4C, D).**

**Fig. 4.**
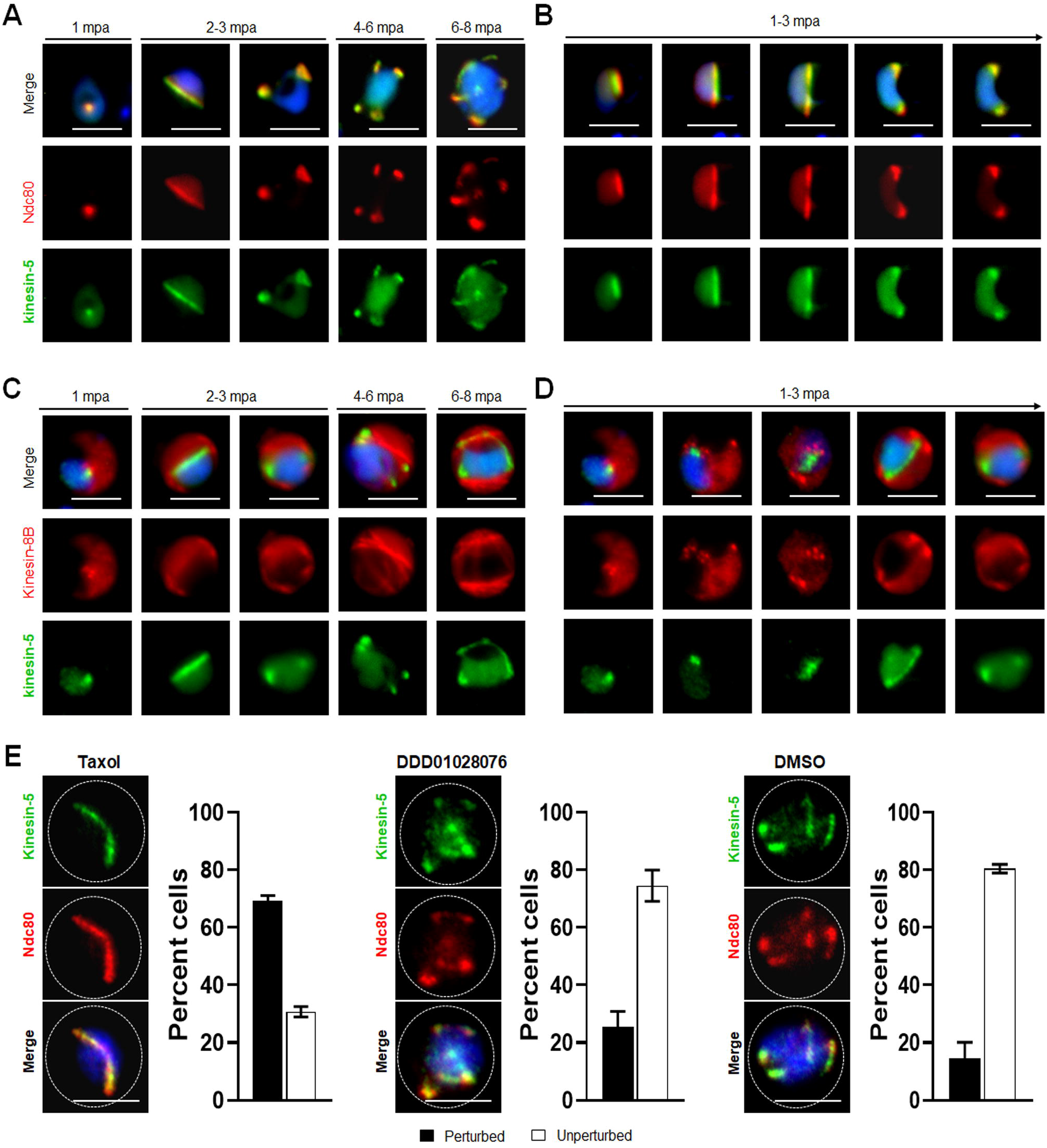
Spatiotemporal dynamics of Pbkinesin-5-GFP with kinetochore marker (NDC80); and basal body and axoneme marker (kinesin-8B), and the effect of MT inhibitors on kinesin-5 distribution. **(A)** The location of kinesin-5-GFP (green) in relation to the kinetochore marker, NDC80-mCherry (red) during male gametogony. (**B)** The dynamic location of kinesin-5-GFP and NDC80-mCherry during the first round of mitosis (1-3 mpa) in male gametogony. (**C)** The location of Pbkinesin-5 in relation to kinesin-8B, a basal body and axonemal marker. The nuclear location of kinesin-5-GFP contrasts with the cytoplasmic location of Pbkinesin-8B during male gametogony. **(D)** The dynamic location of Pbkinesin-5-GFP and kinesin-8B-mCherry during the first round of mitosis (1-3 mpa.) in male gametogony. **(E)** The MT-stabilizing drug Taxol blocks the dynamic distribution of kinesin-5-GFP and NDC80-mCherry; the resulting phenotype of compound addition at 1 mpa is shown. The antimalarial molecule (DDD010128706) had no significant effect on the dynamic location of kinesin-5-GFP and NDC80-mCherry. Inhibitors were added to gametocytes at 1 mpa and cells were fixed at 8 mpa. Scale bar = 5 μm. Bar diagram shows mean ± SEM. n=3.

To study further the association of kinesin-5 with nuclear or cytoplasmic MTs, we examined the effects of tubulin inhibitors specific to nuclear (taxol) and cytoplasmic (DDD01028076) MTs on kinesin-5 organisation during male gametogony (Zeeshan et al., 2019a). Addition of taxol at 1 mpa blocked the dynamic redistribution of both kinesin-5 and NDC80 in more than 70% of male gametocytes, whereas DMSO-treated gametocytes showed normal mitotic progression and kinesin-5 distribution **(Fig. 4D).** This result showed that kinesin-5 distribution and localization is associated with spindle dynamics, similar to the behaviour of NDC80 (Zeeshan et al., 2020b), and can be blocked by taxol treatment, which binds tubulin and stabilises MTs by preventing depolymerisation **(Fig. 4D).** In contrast, treatment with DDD01028076 had no effect on either kinesin-5 or NDC80 location, consistent with the specificity of this inhibitor for cytoplasmic MT (axonemes) (**Fig. 4D**), as shown previously (Zeeshan et al., 2019a).

### During meiosis in zygote to ookinete development, Pbkinesin-5 location follows spindle dynamics

We studied the location of kinesin-5 in the meiotic stage during zygote differentiation to ookinete over the 24-hour period after fertilisation.

Kinesin-5-GFP fluorescence was initially diffuse within the zygote nucleus, but after 1.5 to 2 hours post fertilization the GFP signal coalesced to a single focal point adjacent to the DNA (**Fig. 5A)**. As ookinete development proceeds, with a small apical protrusion (stage I), the intensity of kinesin-5 increased and by stage II to III it was observed on spindles and more prominently on spindle poles (**Fig. 5A)**. Later, in development stage IV, the polar localization of kinesin-5-GFP was lost and by stage V to VI, it again became diffuse in the nucleus with some less-prominent foci (**Fig. 5A**).

**Fig. 5.**
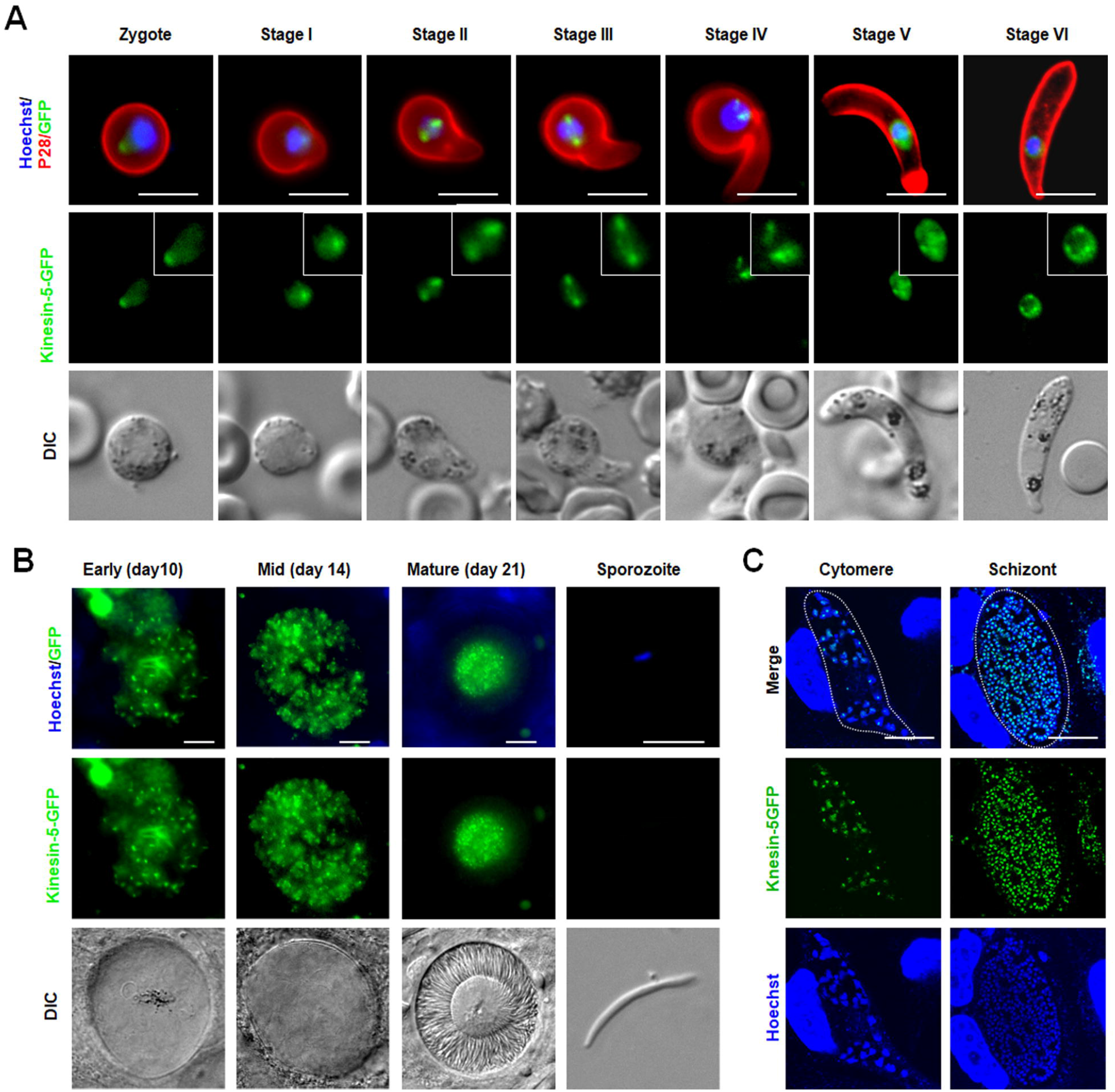
*Pb*kinesin-5 localizes to a spindle and spindle poles during ookinete development, sporogony and liver schizogony. **(A)** Live cell imaging showing *Pb*kinesin-5-GFP location during ookinete development. A cy3-conjugated antibody, 13.1, which recognises the P28 protein on the surface of activated female gametes, zygotes and ookinetes was used to mark these stages (red). Panels: DIC (differential interference contrast), kinesin-5-GFP (green, GFP), Merged: Hoechst (blue, DNA), kinesin-5-GFP (green, GFP) and P28 (red). Scale bar=5 μm. Insets show the zoom of kinesin-5-GFP signal **(B)** Live cell imaging of Pbkinesin-5-GFP in developing oocysts in mosquito guts at 7-, 10-, 14- and 21-days post-infection and in a sporozoite. Panels: DIC (differential interference contrast), kinesin-5-GFP (green, GFP), Merged: Hoechst (blue, DNA) and kinesin-5-GFP (green, GFP). Scale bar = 5 μm **(C)** Expression of the kinesin-5 in early (cytomere) and late liver schizonts detected by live cell imaging. Merge = DAPI and GFP. Dotted lines represent the host cell membrane. Scale bar = 5 μm.

### Pbkinesin-5-GFP exhibits multiple nuclear foci during oocyst development and in liver stage schizogony

Mitosis during oocyst development (sporogony) resembles that of schizogony within the mammalian host, and, as in exoerythrocytic schizogony (in hepatocytes), with many nuclei. Each oocyst contains multiple lobes and produces hundreds of sporozoites (Zeeshan et al., 2020b). Kinesin-5-GFP fluorescence was observed as multiple foci representing a location at the putative MTOC/nuclear poles, from very early in development (day 7) to late stages (at day 14) of oocyst maturation (**Fig. 5B**), similar to the pattern for PbCEN-4 as described previously (Roques et al., 2019). Many arc-shaped GFP signals were also observed that may represent the distribution of kinesin-5 on mitotic spindles (**Fig. 5B**). This mirrors what has been seen in electron microscopy studies: nuclear spindles radiate from the nuclear poles, with attached kinetochores within an intact nuclear membrane during oocyst development (Zeeshan et al., 2020b; Zeeshan et al., 2019b). Interestingly, with maturation of oocyst from day-14 (sporulation started) to day-21 (fully sporulated) kinesin-5 fluorescence started decreasing and in mature oocysts (day 21), fluorescence was restricted to the residual body of the oocyst and was absent from mature sporozoites, suggesting that once nuclear division is completed kinesin-5-GFP is degraded (**Fig. 5B, Fig S3**).

Sporozoites produced in oocysts move to the mosquito salivary glands and, when transmitted by mosquito bite, infect the new host and migrate to the liver. As a model to study expression and location of kinesin-5 during mitosis in liver cells, we infected HeLa cells with sporozoites in vitro. The pattern of kinesin-5 distribution in these cells was similar to that of other asexual proliferative stages showing multiple foci of kinesin-5GFP next to DNA staining (**Fig. 5C**).

## Discussion

Spindle apparatus assembly and chromosome segregation are key processes of nuclear division that require forces generated by MT-based motor proteins (Kull and Endow, 2013; Wordeman, 2010). In most eukaryotes kinesin-5 is the major mitotic motor protein that crosslinks anti-parallel spindle MTs and drives bipolar spindle formation (Ferenz et al., 2010; Shirasugi and Sato, 2019). Any defect in kinesin-5 function results in the failure of spindle pole separation and prevents successful nuclear division (Kapoor et al., 2000; Mann and Wadsworth, 2019; Sharp et al., 1999). In the present study we show by deletion of *kinesin-5* that this protein is not essential for either mitotic or meiotic division during *P. berghei* parasite proliferation but it has an important role in production of infectious sporozoites.

The decrease in number of sporozoites was seen in the oocyst, suggesting an important role for kinesin-5 in sporozoite production in oocysts. Kinesins are MT-based motor proteins and it is not evident if Pbkinesin-5 has any role in regulation of MT function. A recent study has shown that in the absence of MT, chromosome segregation in oocyst proceeds but the MT number and size affect the shape and infectivity of Plasmodium sporozoites (Spreng et al., 2019). The reasons for the drastic decrease in sporozoites in the salivary glands are unclear, but it may result from several factors (Graumans et al., 2020). Either sporozoite egress from the oocyst is impaired or released sporozoites may be unable to reach the salivary glands if they are trapped in other tissues or degraded in the haemolymph by phagocytes (Golenda et al., 1990; Hillyer et al., 2007; Raddi et al., 2020). Another reason is that entry of sporozoites into the salivary glands may be affected, and this depends on the interaction of sporozoite surface proteins, mainly circumsporozoite protein (CSP), thrombospondin related anonymous proteins (TRAP), TRAP related protein (TREP) and apical membrane antigen/erythrocyte binding like protein (MAEBL), with salivary gland proteins such as CSP binding protein (CSPBP), salivary gland surface protein (SGS1) and Saglin protein (Combe et al., 2009; Ghosh et al., 2009; Kariu et al., 2002; Wang et al., 2013). Currently, it is not clear whether kinesin-5 has any role in these events of sporozoite egress or movement or infectivity to salivary glands.

Although the number of sporozoites in salivary glands decreased significantly, they were able to infect a new host. This is likely because even in the absence of kinesin-5, the sporozoite inoculum, number of mosquito bites, or the sporozoite infectivity to liver cells are sufficient to cause infection (Aleshnick et al., 2020; Graumans et al., 2020). Thus, more broadly, the absence of kinesin-5 does not completely block the transmission of parasite.

The biology of malaria parasite development is very different from that of many model eukaryotes, from which it is evolutionarily distinct, with different modes of cell division at different stages. Asexual proliferation is by asynchronous closed mitosis to produce a multinucleate coenocyte that undergoes cytokinesis at the end of the cell cycle to produce haploid extracellular progeny. This cell division lacks several classical regulators such as polo-like kinases, group 1 cyclins and many components of the anaphase promoting complex (Guttery et al., 2014; Roques et al., 2015; Solyakov et al., 2011; Tewari et al., 2010; Wall et al., 2018). The classical cell division kinase, cyclin dependent kinase 1 (CDK1) is not essential for cell division in *Plasmodium* (Tewari et al., 2010), but the parasite does possess other divergent and apicomplexan specific kinases that may be important in this process (Fang et al., 2017; Tewari et al., 2010; Ward et al., 2004). The likely unusual regulatory mechanisms of cell division in *Plasmodium* are consistent with the observation that kinesin-5 is non-essential during most stages of the life cycle except sporozoite production, since there may be alternative, non-classical, ways to mediate and regulate mitosis. Alternatively or additionally, other kinesin motors in *Plasmodium* may have a compensatory role in the absence of kinesin-5; this is seen in budding yeast, where the two kinesin-5s can at least partially complement each other (Hoyt et al., 1992; Roof et al., 1992). Although kinesin-5s and kinesin-13s have very different in vitro activities [32], our transcript analysis of *Δkinesin-5* parasites revealed a significant upregulation of kinesin-13 expression suggesting that this is one mechanistic route by which loss of kinesin-5 in spindle apparatus function can be complemented. Our preliminary data show a similar pattern of expression and localization for kinesin-13-GFP (Zeeshan et al, unpublished), consistent with a compensatory role.

Since *Plasmodium* kinesin-5 is not essential for parasite survival but has an important role in sporozoite production, we were intrigued to see its expression and location during the different mitotic/meiotic stages of the parasite life cycle. Spatiotemporal, dynamic localization of the protein during blood stage schizogony revealed that kinesin-5 starts to coalesce adjacent to the nuclear DNA in the late trophozoite stage. This is the time when the centriolar plaque/ spindle pole body appears for the first time before the start of mitosis, and serves as a putative MTOC (Arnot et al., 2011; Roques et al., 2019). The location of kinesin-5 on this putative MTOC was confirmed by co-localization with centrin-2 and α-tubulin, which have been shown to track MTOC (Arnot et al., 2011; Gerald et al., 2011; Roques et al., 2019). With the progression of nuclear division during schizogony, kinesin-5 showed its characteristic location on spindles, similar to the situation in other eukaryotes, where it helps in spindle assembly and chromosome segregation (Arnot et al., 2011; Ferenz et al., 2010). Kinesin-5 has the same pattern of localization as α-tubulin, showing its association with spindles and MTOCs in consecutive nuclear divisions (Arnot et al., 2011; Ferenz et al., 2010). A similar location of kinesin-5 was observed during the other asexual mitotic stages: liver schizogony and sporogony in the mosquito gut. Kinesin-5 expression was not detected in mature and extracellular merozoites, male gametes and sporozoites, indicating that once its role during mitosis is over it is degraded or discarded. The location of kinesin-5 in the residual body of mature oocysts following release of sporozoites, suggests that kinesin-5 is actively involved during mitosis in oocysts and then it is discarded at the end of endomitotic cell division. A similar fate was also observed for another molecular motor, myosin J (MyoJ) that also accumulates in the residual body during sporogony (Wall et al., 2019).

The residual bodies play important roles in Toxoplasma during organization of developing progeny inside the parasitophorous vacuole and promote their orderly and efficient externalization after maturation (Muniz-Hernandez et al., 2011). A defect in Toxoplasma MyoF molecular motor function results in enlarged residual bodies with accumulation of intact organelles (Jacot et al., 2013), but Pbkinesin-5 deletion did not show any such phenotype. Although the number of *Δkinesin-5* sporozoites in salivary glands is reduced, they are as motile and infective as normal sporozoites, transmitting the parasite.

Plasmodium male gametogony is a very rapid process and completed within 15 minutes, producing eight gametes. It involves three rounds of DNA replication (with 8-fold chromosome replication before nuclear division), along with basal body formation and axoneme assembly in the cytoplasm, followed by chromosome condensation, karyokinesis and cytokinesis leading to the emergence of motile flagellated gametes (Sinden et al., 1976; Zeeshan et al., 2019a). Live cell imaging of kinesin-GFP and fixed immunofluorescence assays using antibodies against GFP and α-tubulin showed that kinesin-5 associates with spindle MTs and spindle poles during the mitotic divisions in male gametogony. The association of kinesin-5 with spindle MTs was further confirmed by gametocyte treatment with Taxol, a spindle MT-specific inhibitor, which inhibited the dynamic relocation of kinesin-5. We investigated further this association of kinesin-5 exclusively with spindle MTs and not with axonemal MTs by live cell imaging of a parasite line expressing both kinesin-5-GFP and kinesin-8B-mCherry. Kinesin-8B is associated with cytoplasmic MTs (axonemes) and not present in the nuclear compartment of male gametocytes (Zeeshan et al., 2019a). Kinesin-5 also associates with kinetochores and this dynamic location of *Plasmodium* kinesin-5 and NDC80 is consistent with a study in yeast, showing that kinesin-5 is recruited to the kinetochore and plays an important role in its organisation (Tytell and Sorger, 2006).

In conclusion, this is the first study to explore the real-time dynamics and functional role of the kinesin-5 molecular motor in the mitotic and meiotic cell division cycles of the different stages of *Plasmodium* development, and to show an important role of kinesin-5 in sporozoite production.

## Material and Methods

### Ethics statement

The animal work performed in this study has passed an ethical review process and was approved by the United Kingdom Home Office. Work was carried out in accordance with the United Kingdom ‘Animals (Scientific Procedures) Act 1986’ for the protection of animals used for experimental purposes under Licence number 40/3344. Six to eight-week-old Tuck’s Original (TO) (Harlan) outbred mice were used for all experiments.

### Generation of transgenic parasites

The gene-deletion targeting vector for *Pbkinesin-5* (PBANKA_0807700) *was* constructed using the pBS-DHFR plasmid, which contains polylinker sites flanking a *T. gondii dhfr/ts* expression cassette conferring resistance to pyrimethamine, as described previously (Tewari et al., 2010). PCR primers N1061 and N1062 were used to generate a 995 bp fragment of *kinesin-5* 5’ upstream sequence from genomic DNA, which was inserted into *ApaI* and *HindIII* restriction sites upstream of the *dhfr/ts* cassette of pBS-DHFR. A 1008 bp fragment generated with primers N1063 and N1064 from the 3’ flanking region of *kinesin-5* was then inserted downstream of the *dhfr/ts* cassette using *Eco*RI and *Xba*I restriction sites. The linear targeting sequence was released using *Apa*I/*Xba*I. A schematic representation of the endogenous *Pbkinesin-5* locus, the construct and the recombined *kinesin-5* locus can be found in **Fig. S1.**

To generate kinesin-5-GFP, a region of *kinesin-5* gene downstream of the ATG start codon was amplified using primers T1921 and T1922 and ligated to p277 vector, and transfected as described previously (Guttery et al., 2012). The p277 vector contains the human *dhfr* cassette, conveying resistance to pyrimethamine. Pbkinesin-5 was tagged with GFP at the C-terminus by single crossover homologous recombination. A schematic representation of the endogenous gene locus, the constructs and the recombined gene locus can be found in **Fig. S2.** The oligonucleotides used to generate the mutant parasite lines can be found in Supplementary **Table S1**. *P. berghei* ANKA line 2.34 (for GFP-tagging) or ANKA line 507cl1 expressing GFP (for gene deletion) parasites were transfected by electroporation (Janse et al., 2006).

### Genotypic analysis of parasites

For the gene knockout parasites, diagnostic PCR was used with primer 1 (IntN106) and primer 2 (ol248) to confirm integration of the targeting construct, and primer 3 (N106 KO1) and primer 4 (N106 KO2) were used to confirm deletion of the *kinesin-5* gene **(Fig. S1)**. For the parasites expressing a C-terminal GFP-tagged kinesin-5 protein, diagnostic PCR was used with primer 1 (IntT192) and primer 2 (ol492) to confirm integration of the GFP targeting construct **(Fig. S2).**

### Phenotypic analyses

To initiate infections, blood containing approximately 60,000 parasites of *Δkinesin-5* line was injected intraperitoneally (i.p.) into mice. Asexual stages and gametocyte production were monitored on Giemsa-stained thin smears. Four to five days post infection, exflagellation and ookinete conversion were examined as described previously (Guttery et al., 2012) with a Zeiss AxioImager M2 microscope (Carl Zeiss, Inc) fitted with an AxioCam ICc1 digital camera. To analyse mosquito transmission, 50-60 *Anopheles stephensi* SD 500 mosquitoes were allowed to feed for 20 min on anaesthetized, infected mice whose asexual parasitemia had reached up to 15% and were carrying comparable numbers of gametocytes as determined on Giemsa-stained blood films. To assess mid-gut infection, approximately 15 guts were dissected from mosquitoes on day 14 post feeding and oocysts were counted on a Zeiss AxioImager M2 microscope using 10x and 63x oil immersion objectives. On day 21 post-feeding, another 20 mosquitoes were dissected, and their guts and salivary glands crushed separately in a loosely fitting homogenizer to release sporozoites, which were then quantified using a haemocytometer or used for imaging and motility assays. Mosquito bite back experiments were performed 21 days post-feeding using naive mice; 15-20 mosquitoes infected with WT-GFP or *Δkinesin-5* parasites were fed for at least 20 min on naive CD1 outbred mice and then infection was monitored after 3 days by examining a blood smear stained with Giemsa’s reagent. For comparison between *Δkinesin-5* and WT-GFP, an unpaired Student’s *t*-test was used.

### Culture and gradient purification of schizonts and gametocytes

Blood cells obtained from infected mice (day 4-5 post infection) were placed in culture for 8-10h and 24 h at 37°C (with rotation at 100 rpm) and schizonts were purified on a 60% v/v NycoDenz (in PBS) gradient, harvested from the interface and washed (NycoDenz stock solution: 27.6% w/v NycoDenz in 5 mM Tris-HCl, pH 7.20, 3 mM KCl, 0.3 mM EDTA). Purification of gametocytes was achieved using a protocol as described previously (Beetsma et al., 1998) with some modifications. Briefly, parasites were injected into phenylhydrazine treated mice and enriched by sulfadiazine treatment after 2 days of infection. The blood was collected on day 4 after infection and gametocyte-infected cells were purified on a 48% v/v NycoDenz (in PBS) gradient. (NycoDenz stock solution: 27.6% w/v NycoDenz in 5 mM Tris-HCl, pH 7.20, 3 mM KCl, 0.3 mM EDTA). The gametocytes were harvested from the interface and washed.

### Live-cell and time-lapse imaging

Different developmental stages of parasite during schizogony, zygote to ookinete transformation and sporogony were analysed for kinesin-5-GFP expression and localization using a 63x oil immersion objective on a Zeiss Axio Imager M2 microscope. Kinesin-5 expression and location was examined during different developmental stages of the parasite life cycle. Purified gametocytes were examined for GFP expression and localization at different time points (0, 1-15 min) after activation in ookinete medium. Images were captured using a 63x oil immersion objective on the same microscope. Time-lapse videos (1 frame every 5 sec for 15-20 cycles) were taken with a 63x objective lens on the same microscope and analysed with the AxioVision 4.8.2 software.

### Generation of dual tagged parasite lines

The kinesin-5-GFP parasites were mixed with either NDC80-cherry or kinesin-8B-cherry parasites in equal numbers and injected into a mouse. Mosquitoes were fed on this mouse 4 to 5 days after infection when gametocyte parasitaemia was high.

These mosquitoes were checked for oocyst development and sporozoite formation at day 14 and day 21 after feeding. Infected mosquitoes were then allowed to feed on naïve mice and after 4 - 5 days these mice were examined for blood stage parasitaemia by microscopy with Giemsa-stained blood smears. In this way, some parasites expressed both kinesin-5-GFP and NDC80-cherry or kinesin-5-GFP and kinesin-8B-cherry in the resultant schizonts and gametocytes, and these were purified, and fluorescence microscopy images were collected as described above.

### Inhibitor studies

Gametocytes from the parasites expressing kinesin-5-GFP and NDC80-cherry were purified as above and treated with Taxol (Sigma) and an antimalarial molecule (DDD01028076) (Zeeshan et al., 2019a) at 1 mpa and then fixed with 4% paraformaldehyde (PFA, Sigma) at 8 min after activation. Dimethyl sulfoxide (DMSO) was used as a control treatment. These fixed gametocytes were then examined on a Zeiss AxioImager M2 microscope.

### Fixed Immunofluorescence Assay

The purified schizonts from PbKinesin-5-GFP parasites were fixed in 2% paraformaldehyde (PFA) and smeared on poly-L-lysine coated slides. Purified gametocytes were activated in ookinete medium then fixed at 1 min, 2 min, 4 min, 6 min, 8 min and 15 min post-activation with 4% PFA diluted in microtubule stabilising buffer (MTSB) for 10-15 min and added to poly-L-lysine coated eight-well slides. Immunocytochemistry was performed using primary GFP-specific rabbit monoclonal antibody (mAb) (Invitrogen-A1122; used at 1:250) and primary mouse anti-α tubulin mAb (Sigma-T9026; used at 1:1000) or mouse anti-centrin mAb (Millipore-04-1624, used at 1:500) or anti-NDC80 (polyclonal sera, a kind gift from Marc-Jan Gubbels). Secondary antibodies were Alexa 488 conjugated anti-mouse IgG (Invitrogen-A11004) and Alexa 568 conjugated anti-rabbit IgG (Invitrogen-A11034) (used at 1 in 1000). The slides were then mounted in Vectashield 19 with DAPI (Vector Labs) for fluorescence microscopy. Parasites were visualised on a Zeiss AxioImager M2 microscope. Pearson’s colocalization coefficient (R) was calculated using image J software (version 1.44).

### Sporozoite motility assays

Sporozoites were isolated from salivary glands of mosquitoes infected with WT-GFP and *Δkinesin-5* parasites on day 21 post infection. Isolated sporozoites in RPMI 1640 containing 3% bovine serum albumin (Fisher Scientific) were pelleted (5 min, 5,000 rpm, 4°C) and used for motility assays as described previously (Wall et al., 2019). Briefly, a drop (6 μl) of sporozoites was transferred onto a microscope glass slide with a cover slip. Time-lapse videos of sporozoites (one frame every 1 s for 100 cycles) were taken using the differential interference contrast settings with a 63x objective lens on a Zeiss AxioImager M2 microscope and analysed with the AxioVision 4.8.2 software. The motility was also analysed by using Matrigel. A small volume (20 μl) of sporozoites, isolated as above was mixed with Matrigel (Corning). The mixture (6 μl) was transferred on a microscope slide with a cover slip and sealed with nail polish. After identifying a field containing sporozoites, time-lapse videos (one frame every 2 s for 100 cycles) were taken using the differential interference contrast settings with a 63x objective lens.

### Liver stage parasite imaging

For *P. berghei* liver stage parasites, 100,000 HeLa cells were seeded in glass-bottomed imaging dishes. Salivary glands of female *A. stephensi* mosquitoes infected with kinesin-5-GFP parasites were isolated and disrupted using a pestle to release sporozoites, which were pipetted gently onto the seeded HeLa cells and incubated at 37 °C in 5% CO2 in complete minimum Eagle’s medium containing 2.5 μg/ml amphotericin B (PAA). Medium was changed 3 hrs after initial infection and once a day thereafter. For live cell imaging, Hoechst 33342 (Molecular Probes) was added to a final concentration of 1 μg/ml, and parasites were imaged at 24, 48, 55 hrs post-infection using a Leica TCS SP8 confocal microscope with the HC PL APO 63x/1.40 oil objective and the Leica Application Suite X software

### qRT-PCR analysis

RNA was isolated from different stages of parasites including all asexual stages, schizonts, gametocytes, ookinete and sporozoites using an RNA purification kit (Stratagene). cDNA was synthesised using an RNA-to-cDNA kit (Applied Biosystems). Gene expression was quantified from 80 ng of total RNA using SYBR green fast master mix kit (Applied Biosystems). All of the primers were designed using primer3 (Primer-blast, NCBI), and amplified a region of 150-200 bp. Analysis was conducted using an Applied Biosystems 7500 fast machine with the following cycling conditions: 95°C for 20 sec followed by 40 cycles of 95°C for 3 sec; 60°C for 30 sec. Three technical replicates and three biological replicates were performed for each assayed gene. The *hsp70* (PBANKA_081890) and *arginyl-t RNA synthetase* (PBANKA_143420) genes were used as endogenous control reference genes. The primers used for qPCR can be found in Supplementary Table1.

### Statistical analysis

All statistical analyses were performed using GraphPad Prism 5 (GraphPad Software). For qRT-PCR, an unpaired t-test was conducted to examine significant differences between wild-type and mutant strains.

## Acknowledgments

We thank Julie Rodgers for helping to maintain the insectary and other technical works. We also thank Professor Marc-Jan Gubbels from Boston College, MA, USA for a kind gift of NDC80 anti-sera. This manuscript has been released as a pre-print at bioRxiv (Zeeshan et al., 2020a).

## Supplementary Materials

**Fig. S1. Generation and genotype analysis of *Δkinesin-5* parasites**

**(A) Schematic of *Plasmodium berghei* kinesin-5 protein (1-1440 aa) showing different domains. Approximate length of different domains is indicated. aa; amino acids. (B)** Schematic representation of the endogenous *kinesin-5* locus, the targeting gene deletion construct and the recombined kinesin-5 locus following double homologous recombination. **(B)** Integration PCR showing correct integration with expected size of bands and deletion of kinesin-5 gene from knockout (mut).

**Fig. S2. Generation and genotypic analysis of kinesin-5-GFP parasites**

**(A)** Schematic representation for 3’-tagging of *kinesin-5* gene with green fluorescent protein (GFP) sequence via single homologous recombination. **(B)** Integration PCR showing correct integration of tagging construct. **(C)** Western blot showing expected size of kinesin-5-GFP protein.

**Fig. S3. Expression and localization of kinesin-5 during sporulation in oocysts**

Live cell imaging showing the kinesin-5-GFP fluorescence in a developing oocyst between day 14 (sporulation starts) and day 21 (completely sporulated) after infection.

**Table S1. Oligonucleotides used in this study**

**Movie S1: Gliding motility of WT-GFP salivary gland sporozoite**

**Movie S2: Gliding motility *Δkinesin-5* salivary gland sporozoite**

**Movie S3: Gliding motility of WT-GFP salivary gland sporozoite on matrigel**

**Movie S4: Gliding motility *Δkinesin-5* salivary gland sporozoite on matrigel**

