## Supplementary figures and images for "*Plasmodium berghei* kinesin-5 associates with the spindle apparatus during cell division and is important for efficient production of infectious sporozoites"

### Supplementar figures

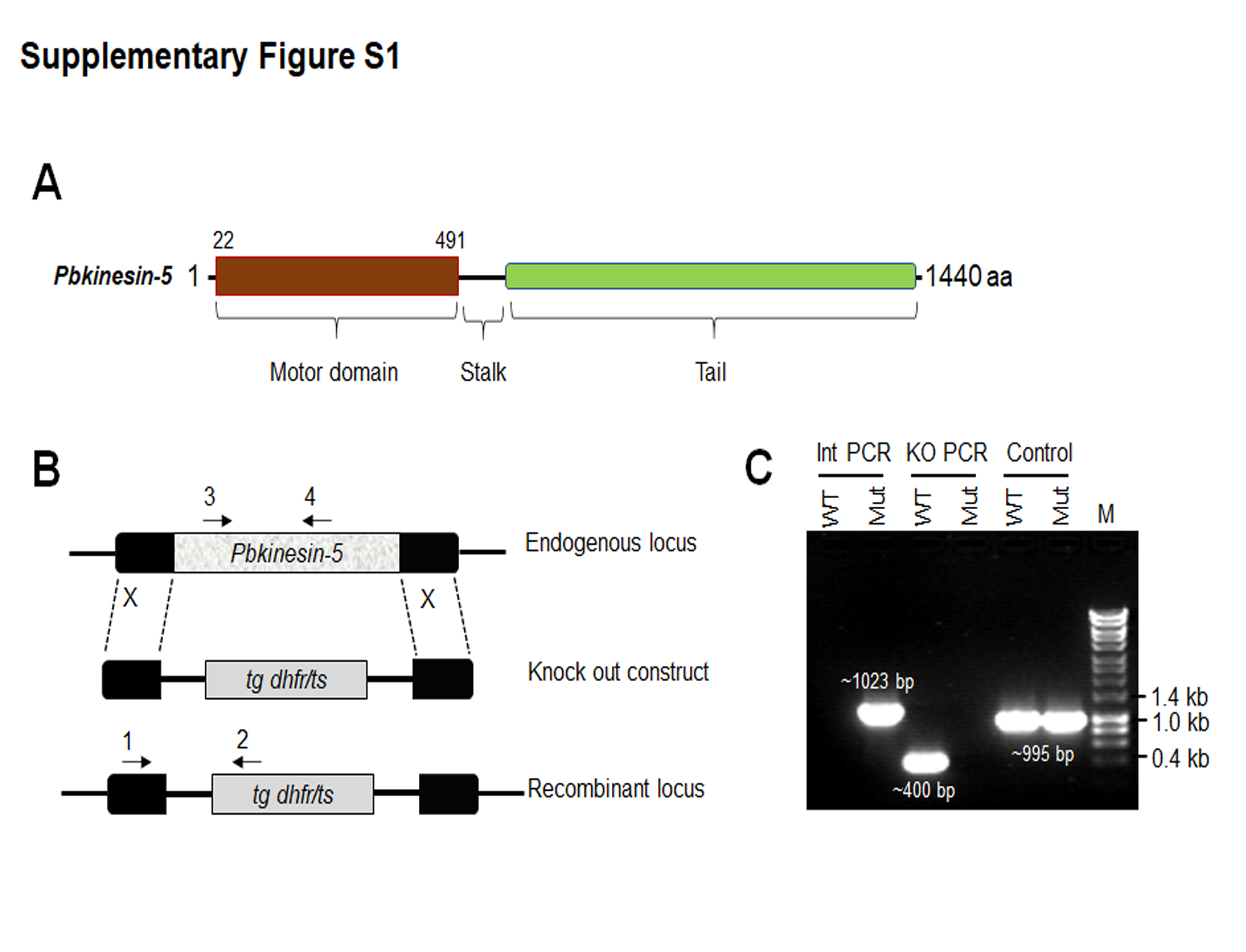

### Supplementary Figure

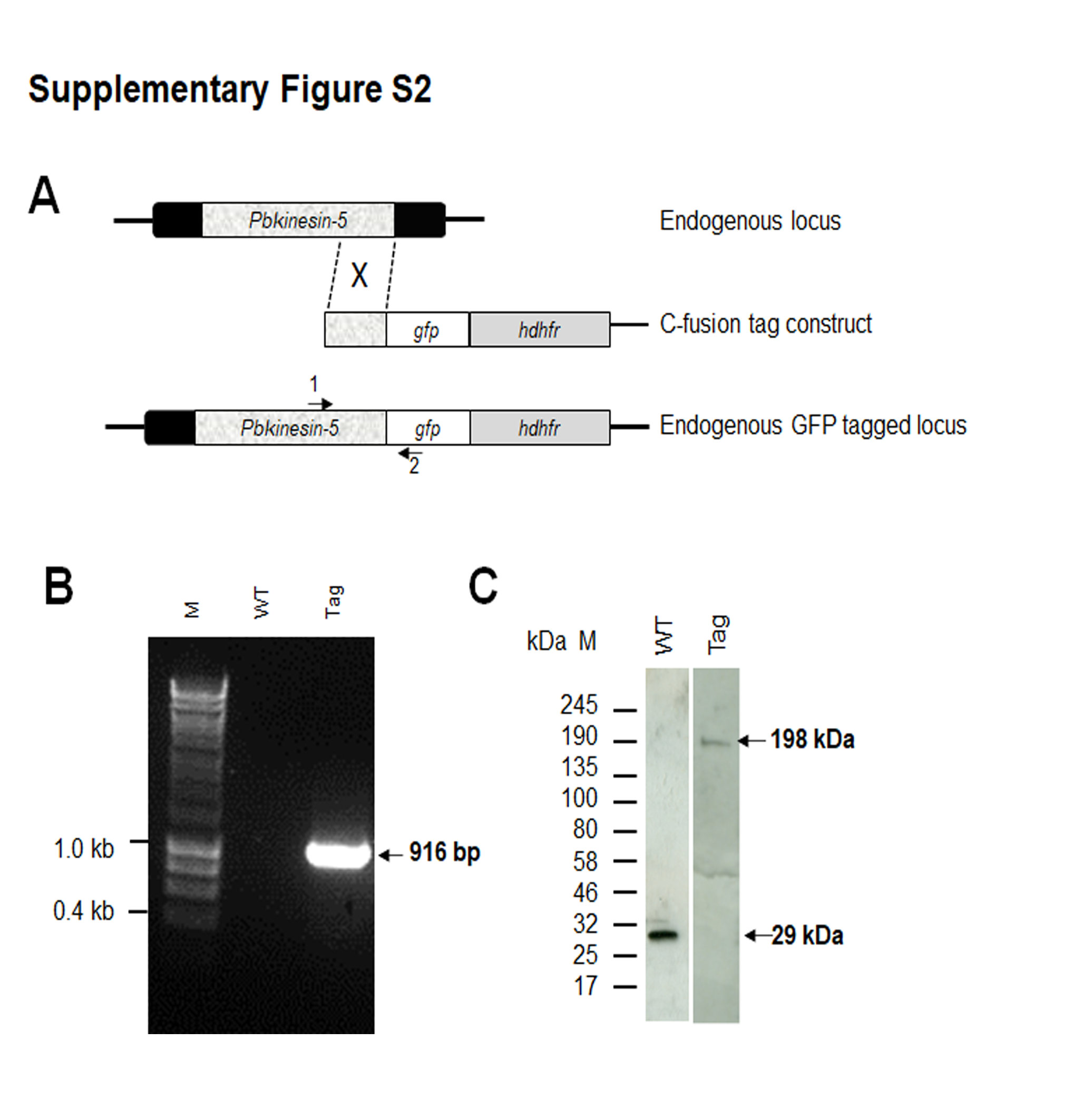

### Supplementary Figure

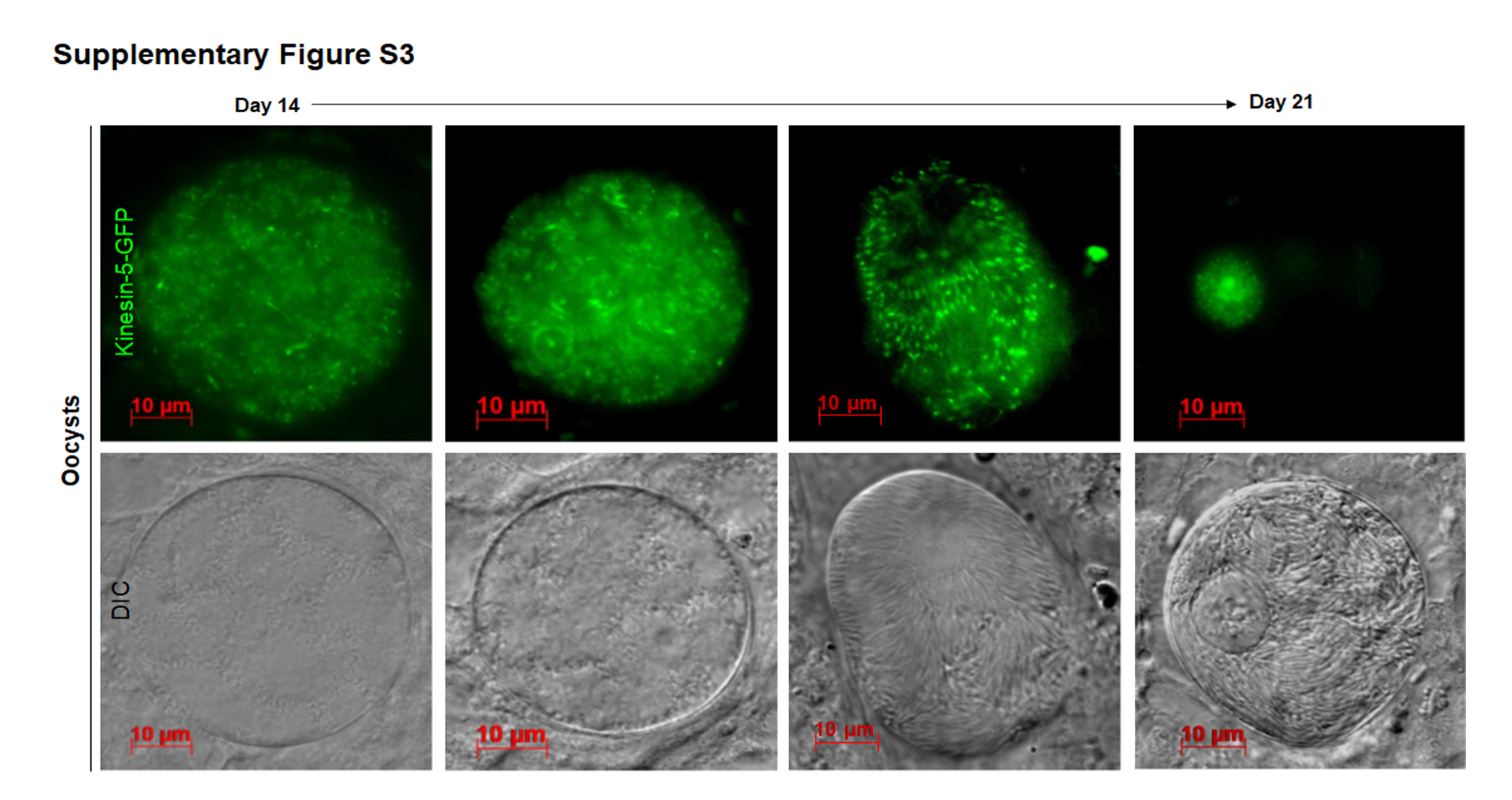
